# Biology and engineering of integrative and conjugative elements: Construction and analyses of hybrid ICEs reveal element functions that affect species-specific efficiencies

**DOI:** 10.1101/2021.12.16.473081

**Authors:** Emily L. Bean, Calvin Herman, Alan D. Grossman

**Affiliations:** Department of Biology, Massachusetts Institute of Technology, Cambridge, Massachusetts 02139

**Author notes:** Corresponding author: Department of Biology, Building 68-530, Massachusetts Institute of Technology, Cambridge, MA 02139.

## Abstract

Integrative and conjugative elements (ICEs) are mobile genetic elements that reside in a bacterial host chromosome and are prominent drivers of bacterial evolution. They are also powerful tools for genetic analyses and engineering. Transfer of an ICE to a new host involves many steps, including excision from the chromosome, DNA processing and replication, transfer across the envelope of the donor and recipient, processing of the DNA, and eventual integration into the chromosome of the new host (now a stable transconjugant). Interactions between an ICE and its hosts throughout the life cycle likely influence the efficiencies of acquisition by new hosts. Here, we investigated how different functional modules of two ICEs, Tn*916* and ICE*Bs1*, affect the transfer efficiencies into different host bacteria. We constructed hybrid elements that utilize the high-efficiency regulatory and excision modules of ICE*Bs1* and the conjugation genes of Tn*916*. These elements produced more transconjugants than Tn*916*, likely due to increased excision frequencies. We also found that several Tn*916* and ICE*Bs1* components can substitute for one other. Using *B. subtilis* donors and three *Enterococcus* species as recipients, we found that different hybrid elements were more readily acquired by some species than others, demonstrating species-specific interactions in steps of the ICE life cycle. This work demonstrates that hybrid elements utilizing the efficient regulatory functions of ICE*Bs1* can be built to enable efficient transfer into and engineering of a variety of other species.

**Author summary (non-technical):** Horizontal gene transfer helps drive microbial evolution, enabling bacteria to rapidly acquire new genes and traits. Integrative and conjugative elements (ICEs) are mobile genetic elements that reside in a bacterial host chromosome and are prominent drivers of horizontal gene transfer. They are also powerful tools for genetic analyses and engineering. Some ICEs carry genes that confer obvious properties to host bacteria, including antibiotic resistances, symbiosis, and pathogenesis. When activated, an ICE-encoded machine is made that can transfer the element to other cells, where it then integrates into the chromosome of the new host. Specific ICEs transfer more effectively into some bacterial species compared to others, yet little is known about the determinants of the efficiencies and specificity of acquisition by different bacterial species. We made and utilized hybrid ICEs, composed of parts of two different elements, to investigate determinants of transfer efficiencies. Our findings demonstrate that there are species-specific interactions that help determine efficiencies of stable acquisition, and that this explains, in part, the efficiencies of different ICEs. These hybrid elements are also useful in genetic engineering and synthetic biology to move genes and pathways into different bacterial species with greater efficiencies than can be achieved with naturally occurring ICEs.

## Introduction

Integrative and conjugative elements (ICEs), also called conjugative transposons, are mobile genetic elements that are major drivers of bacterial evolution (Bellanger et al., 2014; Delavat et al., 2017; Johnson and Grossman, 2015; Roberts and Mullany, 2009; Wozniak and Waldor, 2010). They reside in a host chromosome and can transfer to a recipient cell in a contact-dependent process termed conjugation. ICEs often contain cargo genes that are not needed for the ICE life cycle, but that confer various phenotypes to host cells, including antibiotic resistances, pathogenicity, symbiosis, and various metabolic capabilities (Frost et al., 2005; Johnson and Grossman, 2015; Treangen and Rocha, 2011). In addition to their natural functions, ICEs have been engineered to allow genetic manipulation of a range of organisms (Brophy et al., 2018; Miyazaki and van der Meer, 2013; Peters et al., 2019).

ICEs spend most of their time integrated in and passively propagated with the chromosome of the host cell. Either stochastically or in response to some signal(s), ICEs use element-encoded site-specific recombination machinery to excise from the chromosome to form a plasmid. At this stage, many (perhaps all) ICEs replicate autonomously by rolling-circle replication (Carraro and Burrus, 2015; Delavat et al., 2017; Lee et al., 2010; Thomas et al., 2013; Wright and Grossman, 2016). The element-encoded relaxase nicks and becomes covalently attached to the 5’ end of DNA at the origin of transfer (*oriT*), and host-encoded replication machinery, including a helicase and DNA polymerase, is recruited for unwinding and replicating the ICE DNA (Carraro et al., 2015, 2016; Delavat et al., 2019; Lee et al., 2010; Thomas et al., 2013; Wright and Grossman, 2016).

ICEs encode proteins that comprise the conjugation machinery, a type IV secretion system (T4SS) that transfers DNA from donor to recipient cells. In most cases, the DNA that is transferred is linear, single-stranded, and covalently attached to the relaxase (Draper et al., 2005; Garcillán-Barcia et al., 2007; Llosa et al., 2002). Upon entry into the recipient, the linear ssDNA re-circularizes, becomes double-stranded following replication from a single-strand origin of replication (*sso*) (Wright and Grossman, 2016; Wright et al., 2015), and eventually integrates into the chromosome of its new host to generate a stable transconjugant.

Although most ICEs follow the same general lifecycle, they vary widely in the overall efficiency of transfer. Additionally, different ICEs have different natural host ranges: some ICEs are found in many different bacterial host species, whereas others have been found in only one species. In addition to their natural hosts, many ICEs can be transferred into species in which they are not naturally found (Auchtung et al., 2005; Brophy et al., 2018; Ivins et al., 1988; Miyazaki and van der Meer, 2013; Peters et al., 2019).

The ability of ICE gene products to interact with host components at various steps of the ICE life cycle likely influences the efficiency or effectiveness of transfer out of and into different bacterial hosts. In particular, differences in efficiencies of activation, recombination (excision), DNA processing, unwinding and replication, the compatibility of the conjugation machinery with donor and recipient cell wall and membrane structures, and the availability of a suitable integration site and the ability to integrate in nascent transconjugants could all impact the stable acquisition of an ICE (Delavat et al., 2017; Johnson and Grossman, 2015; Wozniak and Waldor, 2010).

We were interested in determining which step(s) of the ICE life cycle account for differences in conjugation efficiencies into various host species between two well-studied ICEs found in phylogenetically distinct cocci and rod-shaped Gram-positive organisms: Tn*916* and ICE*Bs1*. In addition, because the natural hosts of these elements are different, we thought that insights into which parts of the life cycle led to different efficiencies could allow us to engineer elements with improved function in different hosts.

Tn*916* (∼18 kb) was the first-discovered ICE and carries the tetracycline resistance gene *tetM* (Franke and Clewell, 1981a, 1981b). It and its close relatives are found in *Enterococcus, Streptococcus, Staphylococcus*, and *Clostridium* species (Clewell and Flannagan, 1993; Clewell et al., 1985; Fitzgerald and Clewell, 1985; Roberts and Mullany, 2009, 2011; Sansevere and Robinson, 2017; Santoro et al., 2014). Tn*916* is also functional in *Bacillus subtilis* (e.g. Christie et al., 1987; Mullany et al., 1990; Scott et al., 1988; Wright and Grossman, 2016) and other *Bacillus* species (e.g. Ivins et al., 1988; Naglich and Andrews, 1988).

ICE*Bs1* (∼20.5 kb) of *B. subtilis* encodes conjugation and replication machinery homologous to that of Tn*916*. ICE*Bs1* has only naturally been observed in *B. subtilis*, although related sequences are found in several other *Bacillus* species (Avello et al., 2019; Davis and Grossman, 2021). ICE*Bs1* transfers into a variety of other *Bacillus* species (including *B. anthracis, B. cereus, B. thuringiensis*) and also a variety of *Enterococcus, Streptococcus*, and *Listeria* species (Auchtung et al., 2005; Brophy et al., 2018; DeWitt and Grossman, 2014).

Tn*916* and ICE*Bs1* appear to transfer with different efficiencies, but no direct comparisons have been made. Here, we compare directly the transfer efficiencies of these two elements. We constructed and analyzed hybrid elements that contain the regulatory and integration-excision systems of ICE*Bs1* and the DNA processing and conjugation genes from Tn*916*. We used these hybrid elements to analyze steps in the ICE life cycle that contribute to different conjugation efficiencies. We also used these hybrids to demonstrate the utility of such elements for genetic engineering as these hybrid elements can be acquired more efficiently by some recipient species than either parent element. The approaches described here are generalizable to many other elements and provide a platform for using the regulatory and integration-excision components of ICE*Bs1* for studying conjugation systems from other elements.

## Results

### ICE*Bs1* has higher excision and conjugation efficiencies than Tn*916* in *Bacillus subtilis*

We compared directly the excision and conjugation efficiencies of Tn*916* and ICE*Bs1* in *B. subtilis* using typical mating conditions that enable high conjugation efficiencies (Auchtung et al., 2005). The donor (host) strain with Tn*916* (CMJ253) has a single copy of the element integrated in the genome between *yufK* and *yufL* (see Materials and Methods). Tn*916* is regulated, at least in part, by a transcriptional attenuation mechanism that is partially relieved in the presence of tetracycline or other drugs that inhibit translation (Celli and Trieu-Cuot, 1998; Manganelli et al., 1995; Roberts and Mullany, 2009; Scornec et al., 2017; Showsh and Andrews, 1992; Su et al., 1992). The donor (host) strain with ICE*Bs1* (JMA168), has the element (with an antibiotic resistance gene for selection; see Materials and Methods) integrated in its normal genomic attachment site (*trnS-leu2*). In addition, ICE*Bs1* donor strains contain an inducible copy of the activator *rapI* (Pspank(hy)-*rapI*). Expression of *rapI* causes activation of ICE*Bs1* gene expression and excision, typically in 25-90% of the cells in a population (Auchtung et al., 2005, 2007; Lee et al., 2007). We employed the inducible *rapI* system of ICE*Bs1* because that is what is most useful to achieve high efficiencies of activation in donor cells and as described below, serves as the basis for regulation of hybrid elements.

Strains containing either Tn*916* or ICE*Bs1* were grown to exponential phase in LB medium and activation of each element was stimulated by adding either 2.5 µg/ml of tetracycline or 1mM IPTG for Tn*916* and ICE*Bs1*, respectively, for one hour. We used qPCR to measure excision from the respective integration sites and normalized to a nearby chromosomal locus (*yddN* for ICE*Bs1*, and *mrpG* for Tn*916*) (Lee et al., 2007; Wright and Grossman, 2016). Typically, Tn*916* and ICE*Bs1* had excised from the chromosome in approximately 1-2% and 40% of cells, respectively (Table 1).

**Table 1.** Excision frequencies and conjugation efficiencies of ICEs.

| Donor<sup>a</sup> | Conjugation Efficiency (%) (total donors)<sup>b</sup> | Donor excision frequency (%)<sup>c</sup> | Normalized Conjugation Efficiency (%) (activated donors)<sup>d</sup> |
| --- | --- | --- | --- |
| *Tn916* | 0.0015 ± 0.0002 | 1.5 ± 0.1 | 0.11 ± 0.01 |
| *ICEBs1* (Δ*rapIphrI*) | 2.7 ± 0.2 | 42 ± 4 | 6.7 ± 0.4 |
| *ICEBs1* (Δ*yddJ-yddM*) | 3.2 ± 0.1 | 41 ± 4 | 8.3 ± 0.5 |
| (*ICEBs1*-*Tn916*)-H1<br>(*Tn916* DNA processing) | 0.47 ± 0.06 | 41 ± 2 | 1.2 ± 0.2 |
| (*ICEBs1*-*Tn916*)-H2<br>(*ICEBs1* DNA processing) | 2.0 ± 0.2 | 42 ± 4 | 5.3 ± 0.4 |

<sup>a</sup>Donor strains contained the indicated ICEs: CMJ253 (*Tn916*), JMA168 (*ICEBs1* ( $\Delta$ *rapIphrI*), ELC1211 (*ICEBs1* ( $\Delta$ (*yddJ-yddM*))), ELC1213 (H1), and ELC1185 (H2). *ICEBs1*, H1, and H2 donors also contained *amyE::*[(Pspank(hy)-*rapI*) *spc*] for IPTG-inducible overproduction of *RapI* to stimulate element activation.
<sup>b</sup>Donor strains were grown to exponential phase in LB medium, ICE excision was stimulated with IPTG or tetracycline as appropriate for one hour, before mixing with a recipient strain (CAL419: ICE-cured, *comK::cat*, *str*), filtering, and placing on a solid surface for one hour. Conjugation efficiencies were calculated as the number of transconjugants (*strR* and *kanR* for *ICEBs1*, H1, and H2 matings; *strR* and *tetR* for *Tn916* matings) normalized to the total number of input donors.
<sup>c</sup>Following element induction, DNA was harvested from an aliquot of donor cells. qPCR was used to quantify the amount of empty ICE attachment site relative to a nearby chromosomal locus (see Materials and Methods). The frequency was indicative of the percentage of “activated” donor cells.
<sup>d</sup>The calculated conjugation efficiency was divided by the excision frequency to determine the conjugation efficiency per “activated” donor cells present at the start of the mating. All values are means from 4 or more independent mating assays ± SEM.

Conjugation frequencies were measured by mixing the donor cells containing either Tn*916* or ICE*Bs1* with recipients (devoid of any conjugative element), filtering, and incubating the filters on a solid surface for one hour before selecting for transconjugants (Materials and Methods). Conjugation efficiencies of each element were calculated by normalizing the number of transconjugants to the number of total donor cells (Table 1). Tn*916* and ICE*Bs1* produced approximately 0.0015% and 2.7% transconjugants per donor, respectively, an approximately 1,000-fold difference.

As noted above, there were different frequencies of excision of each element. Because only the excised circular form of either element is competent for transfer, we normalized the number of transconjugants to the excision frequency of each element. After normalization, we found that the conjugation efficiencies of Tn*916* and ICE*Bs1* were 0.11% and 6.7%, respectively, per donor with an excised element. Based on this analysis, we conclude that acquisition of ICE*Bs1* by recipient cells during mating was approximately 50-fold more efficient than that of Tn*916*. This could be due to differences in any of several steps of the ICE life cycle, including DNA nicking, unwinding, replication, association with the conjugation machinery, transfer, second strand DNA synthesis in the recipients, and integration. Based on these considerations, we decided to build a hybrid ICE that uses the regulation, excision, and integration components of ICE*Bs1* and the DNA processing and conjugation components of Tn*916*.

### Design and function of a hybrid conjugative element

Like many ICEs, both Tn*916* and ICE*Bs1* have a modular organization (Burrus and Waldor, 2004; Toussaint and Merlin, 2002), with genes involved in different parts of the life cycle clustered (Figure 1). Because of this modularity, it was relatively straightforward to use the regulatory architecture of ICE*Bs1* (the genes and sequences at the left and right ends) and to replace the DNA processing and conjugation genes with those from Tn*916* (Figure 1c). Such a construct leaves intact the ICE*Bs1* genes that are required for regulation (*immA, immR*), and recombination (*int, xis*). In addition, key DNA sites are also present, including the promoter P*xis* that drives transcription of *xis* and genes downstream, and the left and right ends of the element that contain the recombination sites (*attL* and *attR*). Several genes (*yddJ, spbK*, and *yddM*) near the right attachment site are not required for ICE*Bs1* transfer and were omitted from the hybrid element (Figure 1c). The hybrid element, (ICE*Bs1*-Tn*916*)-H1, or H1 for short, contains a kanamycin resistance gene (*kan*) and H1 is integrated in the genome at the normal ICE*Bs1* attachment site in *trnS-leu2* (Figure 1c).

**Figure 1.**
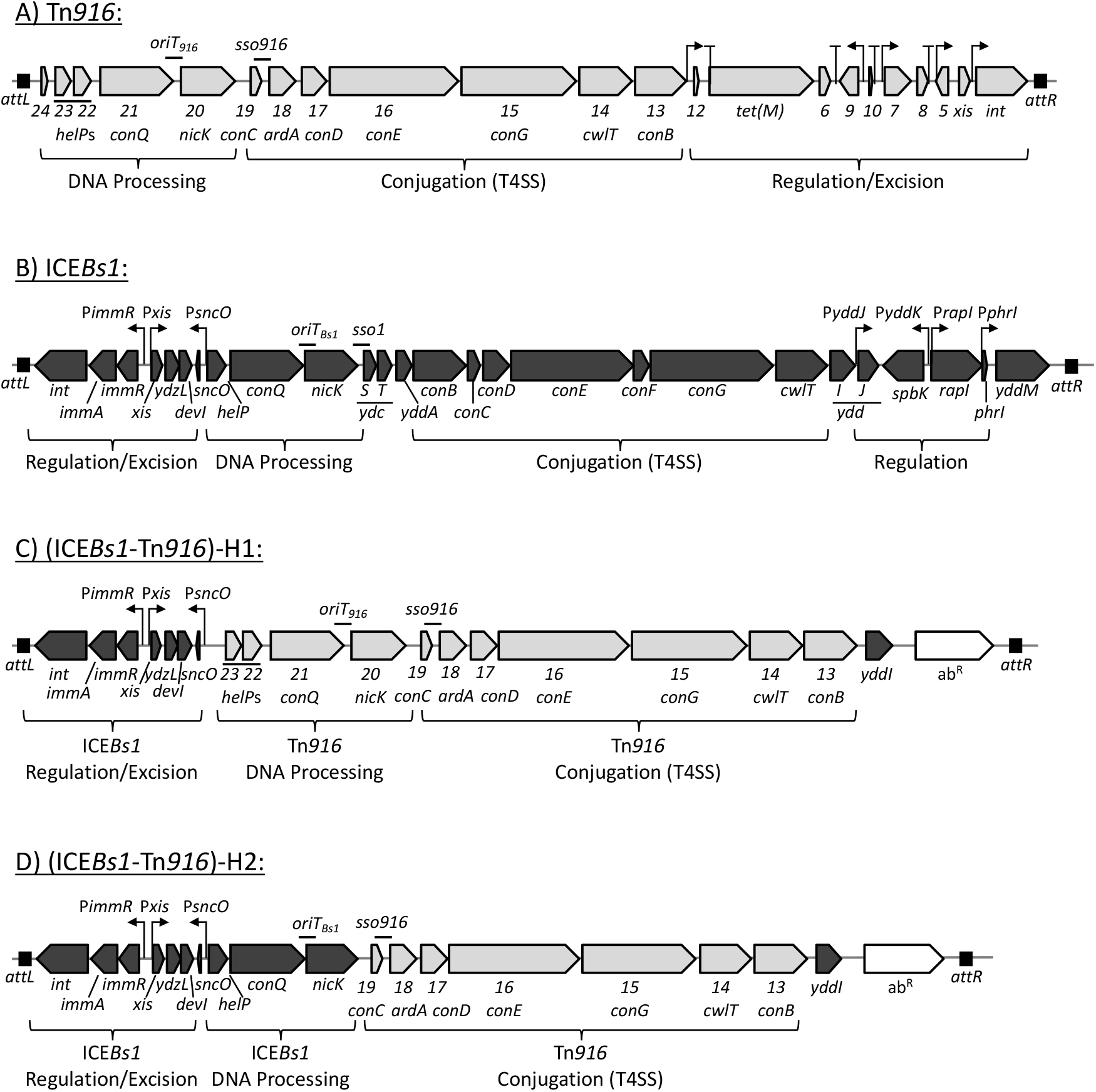
Genetic maps of ICE*Bs1*, Tn*916*, and hybrid elements. Maps are shown of the ICEs used in these experiments: A) Tn*916*; B) ICE*Bs1*; C) (ICE*Bs1*-Tn*916*)-H1 D) (ICE*Bs1*-Tn*916*)-H2. Attachment sites *attL* and *attR* are indicated by black boxes (ICE*Bs1*, (ICE*Bs1*-Tn*916*)-H1), and (ICE*Bs1*-Tn*916*)-H2 are all integrated at *trnS-leu2*; Tn*916* is integrated between *yufK* and *yufL* in donor cells. Open reading frames are indicated by horizontal arrows, pointing in the direction of transcription (gray for Tn*916*, black for ICE*Bs1*). Gene names are located below the depicted open reading frame. Tn*916* gene names are abbreviated to include only the number designation from the gene name (i.e., “*orf23”* is written as “*23*”), and the corresponding ICE*Bs1* homolog gene name is written in parentheses below, when appropriate. *ardA* of Tn*916* encodes an antirestriction protein (Serfiotis-Mitsa et al., 2008). Confirmed and putative promoters are indicated by bent arrows, putative transcription terminators in Tn*916* are indicated by “T” shapes. The current model of transcriptional regulation of Tn*916* (A) is adapted from (Roberts and Mullany, 2009). Previously determined origins of transfer (*oriT*) and single strand origins of replication (*sso*) are indicated by a “-” above the genetic map (Jaworski and Clewell, 1995; Lee and Grossman, 2007; Wright and Grossman, 2016; Wright et al., 2015).

We found that cells containing (ICE*Bs1*-Tn*916*)-H1 (ELC1213) exhibited ICE excision frequencies similar to those containing ICE*Bs1*. Cells were grown in LB medium to exponential phase. Element excision was induced through the addition of 1mM IPTG for one hour during exponential growth. At this time, both ICE*Bs1* and H1 had excised in ∼40% of donor cells (Table 1). These results indicate that, as designed, the element H1 is activated at levels similar to ICE*Bs1*.

Under the same mating conditions as described above, H1 expectedly produced more transconjugants/donor (0.5%) than WT Tn*916* (0.0015%), likely due to its increased activation frequency (Table 1). This is consistent with the increased conjugation efficiencies observed for Tn*916* mutants with increased excision frequencies due to mutations upstream of *tetM*, a region critical for Tn*916* regulation (Beabout et al., 2015). However, H1 consistently produced approximately 5-fold fewer transconjugants/donor than ICE*Bs1* (2.7%). This result indicates that some step(s) other than excision from the chromosome of the donor and integration into the chromosome of the transconjugant has a different efficiency than that of ICE*Bs1*.

The genes *yddJ, spbK*, and *yddM* from ICE*Bs1* are not present in (ICE*Bs1*-Tn*916*)-H1 and we reasoned they might contribute to the different conjugation efficiencies. We deleted these genes in ICE*Bs1* (*ΔyddJ-yddM::kan*) and compared excision and conjugation efficiencies relative to ICE*Bs1* (Δ*rapIphrI*::*kan*). We found that ICE*Bs1* (*ΔyddJ-yddM*::*kan*) and ICE*Bs1* (Δ*rapIphrI::kan*) behaved similarly (Table 1), indicating that there is little if any effect of *yddJ, spbK*, and *yddM* on excision or conjugation efficiencies, consistent with previous findings (Avello et al., 2019; Johnson et al., 2020; Lee and Grossman, 2007). Together, these results indicate that steps in the ICE life cycle after excision are responsible for the differences between ICE*Bs1* and (ICE*Bs1*-Tn*916*)-H1. In addition, because the recombinase and element ends are the same in ICE*Bs1* and H1, integration in the recipient cannot be causing the observed differences. The difference in the conjugation efficiency of each element indicates that some aspect of the conjugation functions encoded by Tn*916* are less efficient than those encoded by ICE*Bs1*.

### Hybrid elements can combine different functional components required for conjugation

Following excision from the host chromosome, ICEs form circular dsDNA intermediates that are processed into a linear, ssDNA form prior to transfer to a neighboring cell (Wozniak and Waldor, 2010). First, the element-encoded relaxase (Orf20 in Tn*916*; NicK in ICE*Bs1*) nicks the DNA substrate at the origin of transfer (*oriT*) and becomes covalently attached to the 5’ end of the DNA. The host-encoded translocase, PcrA, functions as a helicase to unwind the DNA, with the help of an element-encoded helicase processivity factor (Orf23 and Orf22 in Tn*916*; HelP in ICE*Bs1*) (Auchtung et al., 2016; Lee et al., 2010; Thomas et al., 2013; Wright and Grossman, 2016). This nucleoprotein complex, termed the relaxosome, is recognized by the coupling protein (Orf21 in Tn*916*; ConQ in ICE*Bs1*) and subsequently delivered to the T4SS to be transferred to a neighboring cell (Alvarez-Martinez and Christie, 2009; Iyer et al., 2004; Llosa et al., 2002; Schröder and Lanka, 2005). The DNA processing and coupling protein components are encoded together in a module in each element (Figure 1). Because Tn*916* and ICE*Bs1* encode homologous genes required for DNA processing and conjugation, we tested if these components could be substituted between elements.

We constructed a second hybrid element that is identical to (ICE*Bs1*-Tn*916*)-H1, except it contains the DNA processing module from ICE*Bs1* in place of that from Tn*916*. This element, referred to as (ICE*Bs1*-Tn*916*)-H2, or H2, encodes the relaxase, helicase processivity factor, and coupling protein from ICE*Bs1* (Figure 1). H2 contains the single strand origin of replication from Tn*916* (*sso916*, located downstream of *orf19*) for priming second-strand synthesis during rolling-circle replication (Wright and Grossman, 2016). *sso1* of ICE*Bs1* was not included in H2 (Wright et al., 2015).

We found that cells containing H2 (ELC1185) had excision frequencies of ∼40%, similar to those of ICE*Bs1* and H1 (Table 1). The conjugation efficiency of H2 was ∼2.0% transconjugants per donor (Table 1). This is similar to the efficiencies observed for ICE*Bs1* matings, and ∼5 fold greater than that of H1. These results indicated that the T4SS encoded by Tn*916* can support efficient conjugative transfer between *B. subtilis* cells and that the ICE*Bs1*-encoded coupling protein can successfully interact with the T4SS encoded by Tn*916*. They also indicate that the differences in conjugation efficiency between elements using the Tn*916* and ICE*Bs1* machineries are likely due to the relaxase, helicase processivity factor, and/or the coupling protein.

### ICE*Bs1* and Tn*916* coupling proteins can substitute for each other during conjugation

The functional transfer of (ICE*Bs1*-Tn*916*)-H2 demonstrated that the ICE*Bs1* coupling protein can successfully interact with the T4SS encoded by Tn*916*. Coupling proteins must also interact with the substrate that is transferred. In this hybrid element, the coupling protein was interacting with its cognate substrate, the relaxosome (relaxase, *oriT*, and likely the helicase processivity factor) from ICE*Bs1*. In addition to transferring their own nucleoprotein complexes, both Tn*916* and ICE*Bs1* can recognize and mobilize heterologous plasmid substrates that lack their own conjugation machinery (ICE*Bs1*: pC194, pHP13, and pUB110-based pBS42; Tn*916*: pC194, pUB110, and probably others) (Lee et al., 2012; Naglich and Andrews, 1988; Showsh and Andrews, 1999). Due to the ability to transfer similar substrates, we suspected that Tn*916* and ICE*Bs1* could transfer each other’s relaxosome substrates, and that perhaps the encoded coupling proteins could be substituted for one another within the elements. Indeed, preliminary experiments revealed Tn*916* and ICE*Bs1* recognize each other’s relaxosomes. Therefore, we investigated the interchangeability of the coupling proteins between ICE*Bs1* and Tn*916*.

We found that the coupling proteins of Tn*916* (Orf21) and ICE*Bs1* (ConQ) can interact with their non-cognate T4SS and non-cognate relaxosome substrate. We replaced the gene encoding the coupling protein in Tn*916*, ICE*Bs1*, (ICE*Bs1*-Tn*916*)-H1, and (ICE*Bs1*-Tn*916*)-H2 with the homologous gene (*conQ* or *orf21*) from the other element (Materials and Methods). In this way, each of these elements encoded a non-cognate coupling protein in association with the DNA processing components. Additionally, for Tn*916*, ICE*Bs1*, and (ICE*Bs1*-Tn*916*)-H1 these coupling protein replacements required interactions with the non-cognate T4SS. The H2 coupling protein swap (*conQ::orf21*) produced an element encoding the Tn*916* coupling protein and T4SS from Tn*916* and the DNA processing components from ICE*Bs1* (Table 2).

**Table 2.** Coupling proteins can recognize non-cognate relaxosome.

| Parent Element<sup>a</sup> | Coupling protein replacement<sup>b</sup> | DNA processing genes<sup>c</sup> | T4SS genes<sup>c</sup> | Conjugation Efficiency (%) (total donors)<sup>d</sup> | Conjugation efficiency relative to parent element<sup>e</sup> |
| --- | --- | --- | --- | --- | --- |
| Tn916 | ICEBs1 (*conQ*) | Tn916 | Tn916 | 0.0011 ± 0.0002 | 0.71 ± 0.08 |
| ICEBs1 (*ΔrapIphrI*) | Tn916 (*orf21*) | ICEBs1 | ICEBs1 | 0.99 ± 0.1 | 0.35 ± 0.03 |
| H1 | ICEBs1 (*conQ*) | Tn916 | Tn916 | 0.25 ± 0.1 | 0.47 ± 0.07 |
| H2 | Tn916 (*orf21*) | ICEBs1 | Tn916 | 1.4 ± 0.3 | 0.97 ± 0.1 |

<sup>a</sup>Donor strains containing the indicated ICEs had the encoded coupling proteins (Orf21 for Tn916; ConQ for ICEBs1) swapped out.
<sup>b</sup>The indicated coupling protein replacements were made within each element: Tn916(*Δorf21::conQ*) (ELC809), ICEBs1(*ΔconQ::orf21*) (ELC866), H1(*Δorf21::conQ*) (ELC1584), and H2(*ΔconQ::orf21*) (ELC1450). ICEBs1, H1, and H2 donors also contained *amyE::*[Pspank(hy)-*rapI*] *spc* for IPTG-inducible overproduction of RapI to stimulate element excision (activation).
<sup>c</sup>The DNA processing and T4SS genes were not changed from the parent element. For clarity, the origin of these genes is indicated here.
<sup>d</sup>Donor strains were grown to exponential phase in LB medium, ICE excision was stimulated with IPTG or tetracycline as appropriate for one hour, before mixing with a recipient strain (CAL419: ICE-cured, *comK::cat str*), filtering, and placing on a solid surface for one hour. Conjugation efficiencies was calculated as the number of transconjugants (strR and kanR for ICEBs1, H1, and H2 matings; strR and tetR for Tn916 matings) normalized to the total number of input donors.
<sup>e</sup>These mating assays were conducted in parallel with matings from Table 1. The conjugation efficiencies of the element with the coupling protein swaps from (d) were normalized to the conjugation efficiencies of the parent element (with no coupling protein swap) in each experimental replicate to determine if there was a negative effect of using alternative coupling proteins. All values are means from 3 or more independent mating assays ± SEM.

Cells containing an element with a coupling protein swap (ELC809, ELC866, ELC1584, and ELC1450) and cells containing the parent element were mated with recipient strain CAL419 for one hour, as described above. We found that each element containing a coupling protein swap was able to function for conjugative transfer (Table 2). The transfer frequency of Tn*916* (*orf21*::*conQ*) (ELC809) was similar to that of Tn*916* (Table 2), indicating that the coupling protein from ICE*Bs1* was able to function in the context of Tn*916* almost as well as that from Tn*916*. Similarly, ConQ was functional in the context of H1 (*orf21*::*conQ*) (ELC1584), which contains the same DNA processing and T4SS components as Tn*916*. This element mated with efficiencies ∼0.25%. These results indicate that ConQ can substitute for Orf21, meaning ConQ can both recognize the relaxosome and interact with the T4SS from Tn*916*.

Orf21 also functioned in place of ConQ in ICE*Bs1* and H2 during conjugation. ICE*Bs1* (Δ*conQ::orf21*) (ELC866) transferred with an efficiency ∼1%, averaging an approximately 3-fold decrease from ICE*Bs1* (*conQ*) matings. H2 (Δ*conQ::orf21*) (ELC1450) transferred with an efficiency of ∼1%, which was nearly indistinguishable from that of its parent element in side-by-side comparisons.

Coupling proteins are ATPases from the HerA-FtsK superfamily of ATPases and are responsible for recognizing the transfer substrate and physically delivering it to the rest of the conjugation machinery for export out of the cell. It is note-worthy that these coupling proteins interact with both the non-cognate transfer substrate and non-cognate conjugation machinery. Previous studies determined that some coupling proteins, including TrwB of plasmid R388, can interact with non-cognate conjugation machinery, but not non-cognate transfer substrates (Llosa et al., 2003). It was later determined that many conjugative coupling proteins including TrwB, VirD4 of the canonical *Agrobacterium tumefaciens* system, and PcfC of pCF10, contain a so-called “all-alpha domain” that is responsible for conferring the specificity of substrate recognition to these coupling proteins (Whitaker et al., 2015). However, this domain is not present in Orf21, ConQ, and their relatives (Iyer et al., 2004), and it is not yet known how these coupling proteins recognize substrates for transfer.

### Designing hybrid elements for transfer into *Enterococcus* species

Conjugative elements have the potential to be used as genetic tools to engineer microbes for various purposes (Brophy et al., 2018; Miyazaki and van der Meer, 2013; Peters et al., 2019). However, the overall efficiency of transfer directly impacts the efficacy of this approach for genetic engineering. Although ICE*Bs1* and these ICE*Bs1*-Tn*916* hybrids can transfer with similar efficiencies from *B. subtilis* donors to *B. subtilis* recipients, we predicted that these elements might have differing abilities to interact with host machinery of different species, potentially resulting in different transfer efficiencies. For instance, Tn*916* might function more efficiently in one of its natural hosts (e.g., *Enterococcus faecalis*) than ICE*Bs1*, which is only naturally found in *B. subtilis*.

We measured the conjugation efficiency of these elements from *B. subtilis* donor cells into three different *Enterococcus* species: *E. faecalis* (the first-identified host of Tn*916*), *E. caccae*, and *E. durans*. Because *Enterococci* are naturally kanamycin resistant (Murray, 1990), we replaced the kanamycin resistance gene in ICE*Bs1* and the hybrid elements with *tetM* from Tn*916*. These elements are referred to as ICE*Bs1*-*tetM*, (ICE*Bs1*-Tn*916*)-H1-*tetM* (or H1-*tetM*), and (ICE*Bs1*-Tn*916*)-H2-*tetM* (or H2-*tetM*). Additionally, donor strains contained a D-alanine auxotrophy (*Δalr*::*cat*). Cells bearing this mutation will not grow on or in media without the addition of D-alanine (Wecke et al., 1997) thereby serving as a mechanism for selecting against donors (counter-selecting) in a mating assay without using another antibiotic resistance gene (Brophy et al., 2018).

We found that the D-alanine auxotrophy and changing the antibiotic resistance marker in ICE donors did not affect the overall conjugation efficiencies. We compared the mating efficiencies of these donors to the initial donors (*alr*+, *kan* rather than *tetM* in the elements). Donor cells containing Tn*916* (ELC1566), ICE*Bs1*-*tetM* (ELC1795), H1-*tetM* (ELC1722), or H2-*tetM* (ELC1725) were grown in LB medium containing 200 μg/ml D-alanine and mated with the *B. subtilis* recipient CAL419 under standard mating conditions as described above. Conjugation efficiencies were calculated as the number of transconjugants (tetracycline-resistant, D-alanine prototrophs) normalized to the number of donors. These elements exhibited similar conjugation efficiencies as those described for the *alr+* donors (Table 1) and are shown in Figure 2.

**Figure 2.**
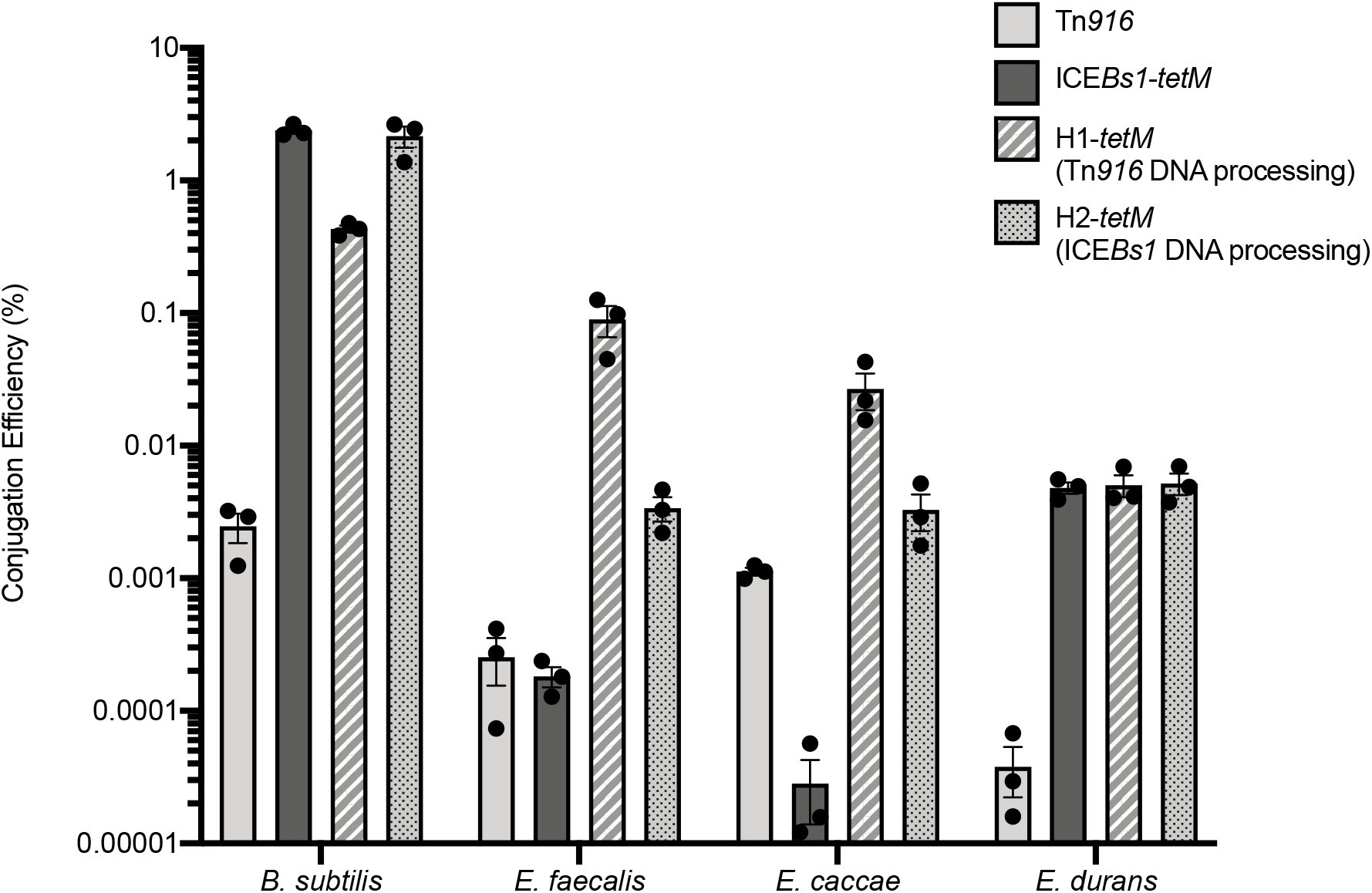
Element conjugation efficiencies are dependent on recipient species. *Bacillus subtilis* donors contained Tn*916* (ELC1566), ICE*Bs1* (ELC1795), (ICE*Bs1*-Tn*916*)-H1 (ELC1722), or (ICE*Bs1*-Tn*916*)-H2 (ELC1725). Donors were D-alanine auxotrophs (*alr::cat*) for counter-selection of transconjugants during mating assays. ICE*Bs1*, H1, and H2 donors also contained *amyE*::[(Pspank(hy)-*rapI*) *spc*] for IPTG-inducible overproduction of RapI to stimulate element excision (activation). Donor cells were grown to exponential phase in LB medium; IPTG or tetracycline was added as appropriate to stimulate element activation. Donors were mixed with recipients that had been grown to exponential phase in LB (*B. subtilis*: CAL419) or BHI (*E. faecalis, E. caccae*, and *E. durans*). Mixed cells were filtered and placed on a solid surface for one hour. Conjugation efficiencies were calculated as the number of transconjugants (tetR, D-alanine prototrophs) produced, normalized to the number of donors. Bars indicated the mean of 3 independent mating assays ± SEM.

### Assessing the species specificity of transfer efficiencies

We found that the conjugation efficiency of the various elements was dependent on the identity of the recipient species. *B. subtilis* donor cells containing Tn*916*, ICE*Bs1*-*tetM*, H1-*tetM*, or H2-*tetM*, were grown in LB medium with D-alanine, as described above. Donor cells were mixed with *E. faecalis* (ATCC 19433), *E. caccae* (BAA-1240), and *E. durans* (ATCC 6056) cells that were grown in BHI medium to exponential phase. Mixtures were filtered and placed on a solid surface for one hour for mating before resuspending cells and selecting for transconjugants that were tetracycline-resistant (ICE+) and D-alanine prototrophs (*Enterococcus*). The conjugation efficiencies were calculated as the number transconjugants produced, normalized to the number of donors applied to the mating. Aliquots of the same donors were used in parallel for all comparisons with different recipients.

When these elements were mated into *E. faecalis* recipients, H1*-tetM*, which utilizes Tn*916* DNA processing and conjugation machinery, consistently produced the most transconjugants/donor (∼0.09%) (Figure 2). H2-*tetM*, which is identical to H1*-tetM* except for its use of the DNA processing module from ICE*Bs1*, produced ∼0.0034% transconjugants/donor, a consistent ∼30-fold decrease compared to H1-*tetM*. These consistent differences indicated that the Tn*916* DNA processing module allows for more efficient acquisition by *E. faecalis* recipients in the context of these hybrid elements. Compared to the hybrid elements, ICE*Bs1* produced the fewest transconjugants per donor (0.00018%), a nearly 20-fold decrease compared to H2*-tetM*. H2-*tetM* differs from ICE*Bs1* through its use of the Tn*916* T4SS and single strand origin (*sso916*), indicating that one, or both, of these Tn*916* features contributed to more efficient acquisition by *E. faecalis*. Notably, Tn*916* produced similar numbers of transconjugants/donor as ICE*Bs1* (0.00025%). This result is in direct contrast to results observed when these elements were mated into *B. subtilis* recipients. The lower activation frequency of Tn*916* (∼0.66 ± 0.08% in donors immediately prior to the start of these matings) compared to that of ICE*Bs1* (∼22 ± 3%) did not correspond to lower conjugation efficiencies, indicating that steps downstream of element activation allowed for Tn*916* to produce more transconjugants.

Similar results were observed when these elements were mated into *E. caccae*, which was first isolated from human stool samples in 2006 (Carvalho et al., 2006). H1-*tetM* produced the most transconjugants/donor (0.027%), which was nearly 10-fold more efficient than H2-*tetM* (0.0033%) (Figure 2). Tn*916* produced more transconjugants/donor (0.0011%) than ICE*Bs1*, which produced the fewest transconjugants of the elements tested (2.8 × 10^−5^ %). Together, these results indicate that both hybrid elements (H1-*tetM*; H2-*tetM*) are more readily acquired by *E. faecalis* and *E. caccae* than either parent element (Tn*916* and ICE*Bs1*). Notably, Tn*916* produced more transconjugants than ICE*Bs1* into these species, indicating there is a benefit to using Tn*916* genes required for conjugation. H1-*tetM* and H2-*tetM* had the advantage in that they utilize components of Tn*916* DNA processing and conjugation machinery, but they activated at higher frequencies in the donor cells (∼20 ± 1%, ∼22 ± 8%, respectively during these experiments) than Tn*916* (∼0.66 ± 0.08%) allowing more mating events to occur.

In contrast, the relative mating efficiencies were quite different with *E. durans* as a recipient. *E. durans* belongs to a distinct phylogenetic *Enterococcus* group (*E. faecium* group) than *E. faecalis* and *E. caccae* (*E. faecalis* group) (Byappanahalli et al., 2012). H1*-tetM*, H2-*tetM*, and ICE*Bs1*-*tetM* all mated with similar efficiencies into *E. durans* (0.0050%, 0.0052%, and 0.0048%, respectively) (Figure 2). In contrast to the *E. faecalis* and *E. caccae* recipient matings, Tn*916* produced the fewest transconjugants/donor (3.8 ×10^−5^ %). These results indicated that Tn*916* and ICE*Bs1* DNA processing and conjugation machinery perform equally well for mating into *E. durans*, with no distinct advantage for either element. Tn*916* likely produced the fewest transconjugants due to its low activation frequency in donor cells.

Altogether, these experiments show that the identity of the recipient species impacts how efficiently a conjugative element can be acquired. Various host factors may impact how efficiently these elements can be received, including the compatibility of cell envelope structures with the conjugation machinery and the ability of these DNA elements to be processed in their new host cells upon acquisition. Furthermore, hosts may be armed to destroy newly-acquired foreign DNA with various defense systems, including restriction modification systems and CRISPR-mediated interference (Bellanger et al., 2014; Cury et al., 2018; Johnson and Grossman, 2015; Serfiotis-Mitsa et al., 2008; Tock and Dryden, 2005; Westra et al., 2012; Wilson and Murray, 1991; Zhang et al., 2013).

### Detecting stable acquisition of conjugative elements in transconjugants

We found that ICE*Bs1* and the hybrid elements did not consistently integrate into the chromosome of the recipient cells, and thus were not stably maintained. We re-streaked transconjugants obtained from these mating assays non-selectively and subsequently checked for tetracycline resistance. We found that fewer than 10% of the apparent transconjugants had maintained the element (Table S1). In the few apparent transconjugants that retained tetracycline resistance, we used arbitrary PCR to map the insertion sites and frequently detected circular ICEs, indicating that the element was not stably integrated into the chromosome (Brophy et al., 2018; Das et al., 2005). We only identified a few stable integration events in *E. caccae* and *E. durans* from H2-*tetM* matings into sites that resembled the preferred attachment site of ICE*Bs1* (Table S1). This element went to a different location in each stable transconjugant, indicating that there is not one preferred attachment site for the element in these chromosomes. We did not identify any integration sites in *E. faecalis*. As previously reported (Brophy et al., 2018), these *Enterococcus* genomes do not contain ideal ICE*Bs1* integration sites (no motifs present with >85% identity with the ICE*Bs1* 17 bp attachment site (Lee et al., 2007)). However, we have previously reported that ICE*Bs1* can integrate into sites within the *B. subtilis* chromosome containing as many as 12 mismatches from the ideal 17 bp attachment site (Menard and Grossman, 2013).

Tn*916* was stably maintained in the transconjugants following non-selective growth. We mapped the integration sites of three transconjugants of each species and identified three unique, frequently intergenic, AT-rich integration sites in each, as expected (Table S1) (Cookson et al., 2011; Mullany et al., 2012; Scott et al., 1994). Because Tn*916* does not integrate in one defined chromosomal site, but instead within an AT-rich region that can be found in many places on a chromosome, Tn*916* had the advantage in target site selection and thus stable acquisition during these matings. For this reason, although Tn*916* produced fewer apparent transconjugants than H1*-tetM* or H2-*tetM*, its attachment site availability can confer an advantage during mating events. By using attachment sites from ICE*Bs1*, the hybrid element is more limited in its ability to integrate into heterologous host chromosomes.

Because the hybrid element containing Tn*916* DNA processing machinery mated more efficiently into *E. faecalis* and *E. caccae* than the hybrid containing ICE*Bs1* DNA processing machinery, we predict that the DNA processing machinery encoded by Tn*916* may be better suited for interaction with these *Enterococcus* species’ host machinery. These interactions with host machinery are necessary for the element to undergo autonomous replication upon acquisition. Because ICE*Bs1* and the hybrid elements frequently did not stably integrate into these recipient chromosomes, improved replication efficiencies could improve the number of detectable transfer events. Unlike ICE*Bs1*, Tn*916* encodes two helicase processivity factors. We have not yet investigated if one or both are required to support element replication, or if these requirements change in the context of different host cells.

## Discussion

Here, we showed how a hybrid element encoding the high-efficiency regulation and integration-excision functions of ICE*Bs1* and the DNA processing and T4SS functions from Tn*916* allowed us to better define the rate-limiting steps of Tn*916* and ICE*Bs1* conjugation from *B. subtilis* hosts into various recipient species that can be phylogenetically distinct. Together, our results indicate that the host range and transfer efficiencies of conjugative elements is dictated by several different stages of the ICE life cycle.

The hybrid elements we generated demonstrate the practicality and efficiency of using the regulation and excision-integration functions of ICE*Bs1* for studying the conjugation machinery and various functions encoded by other conjugative elements. The conjugation machinery of other elements could easily be used in place of that encoded by ICE*Bs1*, just as was the case for that from Tn*916*. The ability to induce ICE*Bs1* by overproduction of the activator RapI should allow investigation of ICEs that may be difficult to study on the population-level due to limited activation. Additionally, the use of such hybrid elements might allow future host range optimization of elements for purposes of genetic engineering. It may be possible to mix and match DNA processing and conjugation machinery from different elements to be compatible with a desired recipient species. However, it is worthwhile to note that although the ICE*Bs1* DNA processing machinery and coupling protein worked with Tn*916* conjugation machinery (in H2), this will not always be the case for other hybrid elements; many elements will likely require the use of cognate transfer substrates and coupling proteins for successful transfer.

In designing future hybrid conjugative elements, it would be feasible to include genes, such as *ardA* of Tn*916* (Serfiotis-Mitsa et al., 2008) to help a hybrid element defend itself against nucleolytic attack by the recipient host cell. Additionally, we imagine it would be possible to install various conjugation machines to allow different transfer substrates, such as effector proteins, to be moved into recipient cells (Bhatty et al., 2013; Segal et al., 1998; Souza et al., 2015). In summary, this system can be used as a chassis for future studies of Tn*916* and a variety of other conjugative elements.

## Materials and Methods

### Media and growth conditions

*B. subtilis* and *Escherichia coli* cells were grown shaking at 37°C in LB medium for all routine growth and strain constructions. *Enterococcus* cells were grown shaking at 37°C in brain heart infusion (BHI) (Becton Dickinson, 237500) medium. When appropriate, cells were grown in liquid culture with 2.5 μg/ml kanamycin, 3 μg/ml tetracycline, or 50 μg/ml spectinomycin, to maintain an ICE or plasmid within a donor cell (cells bearing Tn*916* were not grown in the presence of tetracycline except for periods of induction). Where indicated, donor cells containing ICE*Bs1* or a hybrid element were induced with 1mM isopropyl-β-D-thiogalactoside (IPTG) and donor cells containing Tn*916* were induced with 2.5 μg/ml tetracycline (Auchtung et al., 2005; Wright and Grossman, 2016). Cells with the D-alanine auxotrophy were grown with 200 μg/ml D-alanine. Antibiotic and other supplement concentrations for growth on solid media were: 5 μg/ml kanamycin, 12.5 μg/ml tetracycline, 5 μg/ml chloramphenicol, 100 μg/ml streptomycin, or 400 μg/ml D-alanine as appropriate.

### Strains, alleles, and plasmids

*E. coli* strain AG1111 (MC1061 F’ *lacI*^q^ *lacZM15* Tn*10*) was used for plasmid construction. *Enterococcus* strains were from the ATCC and included: *E. faecalis* ATCC-19433, *E. caccae* ATCC BAA-1240, and *E. durans* ATCC 6056.

*B. subtilis* strains (Table 3), except BS49, were all derived from JH642, contain the *trpC2 pheA1* alleles (Perego et al., 1988; Smith et al., 2014) and were made by natural transformation (Harwood and Cutting, 1990). The construction of various alleles and the hybrid conjugative elements is summarized below.

**Table 3.**
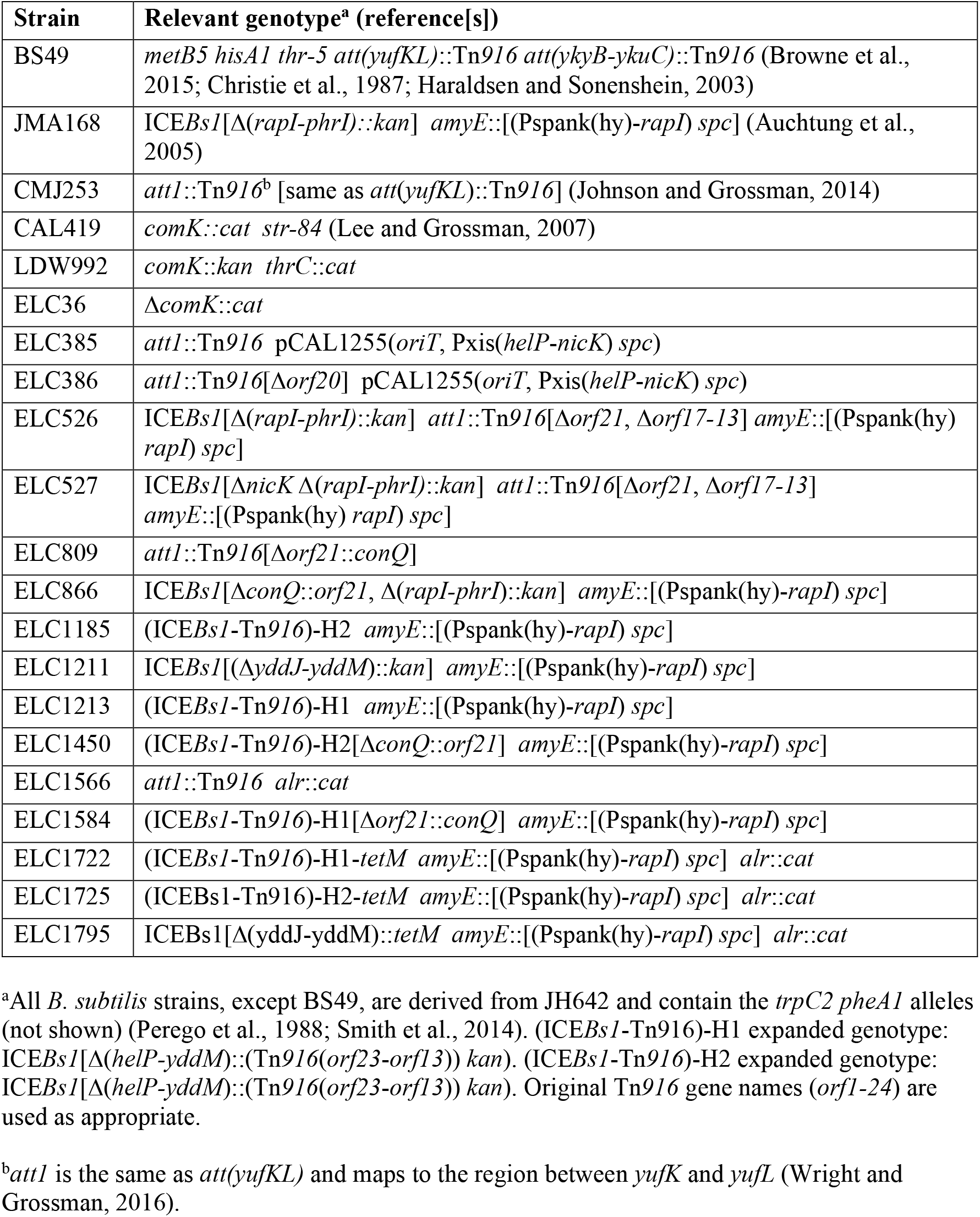
*Bacillus subtilis* strains used.

ICE*Bs1* or hybrid elements were activated by overexpression of *rapI* using an isopropyl-β-D-thiogalactopyranoside (IPTG)-inducible copy of *rapI* (*amyE::*[(Pspank(hy)*-rapI) spc*]) (Auchtung et al., 2005). The Δ(*rapI-phrI*)::*kan* (Auchtung et al., 2005) and Δ*nicK* (Lee and Grossman, 2007) alleles in ICE*Bs1*, the Δ*orf20* (Wright and Grossman, 2016) allele in Tn*916*, and pCAL1255 (Thomas et al., 2013) were all described previously.

The Δ*comK*::*cat* allele in ELC36 replaced most of the *comK* open reading frame from 47bp upstream of *comK* to 19bp upstream of its stop codon with the chloramphenicol resistance cassette from pGEMcat via isothermal assembly of the cassette and up- and downstream homology arms (Gibson et al., 2009).

D-alanine auxotrophs were generated by using linearized pJAB403 to replace the open reading frames of *alrA-ndoA* (using the same borders as previously described (Brophy et al., 2018)) with the chloramphenicol resistance gene from pC194.

### Construction of hybrid elements

(ICE*Bs1*-Tn*916*)-H1 is a large deletion-insertion that removes the ICE*Bs1* DNA processing and conjugation genes from the start codon of *helP* through to 17bp upstream of *yddI* and inserts Tn*916* genes from 121bp upstream of the *orf23* start codon to the stop codon of *orf13*. An additional deletion-insertion removes ICE*Bs1* genes *yddJ*-*yddM* starting 7bp downstream of *yddI*’s stop codon through to the *yddM* stop codon and inserts a kanamycin resistance gene from pGK67 that is codirectional with the genes in the P*xis* operon. These fragments were fused by isothermal assembly (Gibson et al., 2009) with homology arms and transformed into AG174, which contains a copy of ICE*Bs1* at *trnS-leu2*, selecting for acquisition of kanamycin resistance. ICE*Bs1* Δ*yddJ-yddM*::kan is an insertion-deletion of ICE*Bs1* identical to that contained in (ICE*Bs1*-Tn*916*)-H1.

(ICE*Bs1*-Tn*916*)-H2 is an insertion-deletion of the Tn*916* DNA processing machinery in (ICE*Bs1*-Tn*916*)-H1 removing from 121bp upstream of the *orf23* start codon to 42bp upstream of the *orf19* start codon and inserting ICE*Bs1* from the *helP* start codon to the *nicK* stop codon (such that this element was identical to WT ICE*Bs1* from *attL*-*nicK*). Two 1kb DNA fragments containing DNA flanking this region in (ICE*Bs1*-Tn*916*)-H1 were amplified and fused with the insert from ICE*Bs1* and inserted into pCAL1422, a plasmid containing *E. coli lacZ* (Thomas et al., 2013), cut with BamHI and EcoRI, via isothermal assembly to generate pELC1091. The resulting plasmid was integrated by single crossover into a strain containing (ICE*Bs1*-Tn*916*)-H1. Transformants were screened for loss of *lacZ*, indicating loss of the integrated plasmid and checked by PCR for replacement of the DNA processing genes, thereby generating (ICE*Bs1*-Tn*916*)-H2. In generating this functional hybrid element, we determined the Tn*916* gene *orf19*, a predicted integral membrane protein is required for conjugative transfer.

Tetracycline-resistant variants of these ICEs were constructed by replacing *kan* with *tetM* from Tn*916*. The *tetM* insert began 274 bp upstream of *orf12* (*tetM* leader peptide) and ended with the stop codon of *tetM*. This replacement left 105 bp downstream of the *kan* ORF to serve as a transcriptional terminator. Isothermal assembly was used to fuse this replacement antibiotic cassette with ∼1kb of neighboring homology regions before transforming into the appropriate parent strain and selecting for transformants that were tetracycline resistant and confirming kanamycin sensitivity.

A form of Tn*916* lacking all components of its T4SS (Tn*916*[Δ*orf21*, Δ*orf17-13*] was generated through two rounds of deletions. *Δorf21* is an unmarked deletion, removing the first 1272 bp of *orf21* and leaving the remaining 114 bp (and *oriT*) intact. This deletion was amplified with ∼1kb homology arms from pLW805 [*oriT*(*916*), Pspank-*orf20-orf22-orf23*], a plasmid that contains the upstream genes (*orf23, orf22*) fused to the downstream *orf20* (Wright and Grossman, 2016). This amplification was inserted into pCAL1422 cut with BamHI and EcoRI to generate pELC1068. Δ*orf17-orf13* fuses the ninth codon of *orf17* to the 302^nd^ codon of *orf13* within WT Tn*916*. Approximately 1kb homology arms were amplified and inserted into pCAL1422 cut with BamHI and EcoRI via isothermal assembly to generate pELC421. These plasmids were used to make deletions as described above. LDW981 (Tn*916* [Δ*orf21*]) was generated first and then used to generate ELC423 (Tn*916*[Δ*orf21*, Δ*orf17-13*]).

Δ*conQ*::*orf21* is a deletion-insertion that replaces codons 1-373 (of 480 codons) of *conQ* (ICE*Bs1*) with *orf21* (Tn*916*). The last 321 bp of *conQ* were left intact so as not to disrupt *oriT*, as previously described (Lee et al., 2012). Δ*orf21*::*conQ* is a deletion-insertion of *orf21* (of Tn*916*), removing codons 1-424 (of 461 codons) of *orf21* and inserting the entire open reading frame of *conQ* (of ICE*Bs1*). The last 111 bp of *orf21* were left intact so as not to disrupt *oriT*. For either allele, the replacement coupling protein was fused with 1kb DNA amplifications of DNA flanking the originally encoded coupling protein and inserted into the EcoRI and BamHI cut sites of pCAL1422 by isothermal assembly. The resulting plasmids (pELC1389 and pELC801, respectively) were used to generate the deletion-insertions.

### Mating assays

Mating assays were performed essentially as previously described (Auchtung et al., 2005). Briefly, donor cells contained an ICE marked with either a kanamycin or tetracycline antibiotic resistance cassette. Donor cells containing either ICE*Bs1* or a hybrid element were grown in LB medium in the presence of its respective antibiotic (2.5 μg/ml kanamycin or 3 μg/ml tetracycline) to maintain ICE. Donor cells containing Tn*916* were grown non-selectively in LB medium. All donors were grown with D-alanine (200 µg/ml), as needed. *B. subtilis* recipient cells (typically CAL419, unless otherwise indicated) were also grown in LB medium, did not possess any ICE, were resistant to streptomycin (*str-84*), and were defective in competence (*comK*) (Lee and Grossman, 2007). *Enterococcus* recipient cells were grown in BHI medium and were D-alanine prototrophs.

*B. subtilis* donor strains were grown for at least three generations in LB medium to an OD600∼0.2 before stimulating ICE*Bs1* or Tn*916* activation with either 1 mM IPTG or 2.5 μg/ml tetracycline, as appropriate. After 1h, when donor and recipient cultures were at an OD600∼1.0, they were mixed in a 1:1 ratio (5 total ODs of cells) and applied to a nitrocellulose filter for a solid-surface mating. At this point, cells were also harvested for DNA isolation to determine via qPCR the percentage of donor cells with excised ICEs (see below). Filters were incubated on a 1.5% agar plate containing 1X Spizizen salts (Harwood and Cutting, 1990) at 37°C for 1-3h. Cells were harvested off the filters and dilutions were plated on LB or BHI plates containing the appropriate selections to enumerate the number of transconjugants. The conjugation efficiency was calculated as the number of transconjugants present at the end of the mating (CFU/ml) divided by the number of donor cells applied to the mating (CFU/ml) (pre-mating donor counts were used to prevent an overestimation of efficiency due to a drop in viability of donor cells during the course of the mating). Where indicated, mating efficiencies were also normalized to the number of donor cells from which ICE*Bs1* or Tn*916* had excised at the start of the mating (see below).

### Excision assays

We used qPCR to quantify excision of these elements. Genomic DNA of donor cultures was harvested using the Qiagen DNeasy kit with 40 µg/ml lysozyme. The following primers were used to amplify the empty ICE attachment site in the chromosome (only present in cells with excised ICE) and a nearby chromosomal locus for normalization (present in every cell).

For ICE*Bs1* and hybrid elements (integrated at *trnS-leu2*), oMA198 (5’- GCCTACTAAA CCAGCACAAC) and oMA199 (5’- AAGGTGGTTA AACCCTTGG) amplified the empty chromosomal attachment site (*attB*). oMA200 (5’- GCAAGCGATC ACATAAGGTT C) and oMA201 (5’- AGCGGAAATT GCTGCAAAG) amplified a region within the nearby gene, *yddN*.

For Tn*916* (integrated between *yufK* and *yufL*) we used previously described primers (Wright and Grossman, 2016). oLW542 (5’- GCAATGCGAT TAATACAACG ATAC) and oLW543 (5’- TCGAGCATTC CATCATACAT TC) amplified the empty chromosomal attachment site (*att1*). oLW544 (5’- CCTGCTTGGG ATTCTCTTTA TC) and oLW545 (5’- GTCATCTTGC ACACTTCTCT C) amplified a region within the nearby gene *mrpG*.

qPCR was performed using SSoAdvanced SYBR master mix and the CFX96 Touch Real-Time PCR system (Bio-Rad). Excision frequencies were calculated as the number of copies of the empty chromosomal attachment sites (as indicated by the Cp values measured through qPCR) divided by the number of copies of the nearby chromosomal locus. Standard curves for these qPCRs were generated using *B. subtilis* genomic DNA that contained empty ICE attachment sites and a copy of the nearby gene (*yddN* or *mrpG*). DNA for the standard curves was harvested when these strains were in late stationary phase and had an *oriC*/*terC* ratio of ∼1, indicating that the copy numbers of these targets were in ∼1:1 ratios.

### Mapping ICE integration sites

Arbitrary PCR was used to map ICE integration sites, as previously described (Brophy et al., 2018; Das et al., 2005). Following the initial post-mating selection step, transconjugant colonies were re-streaked non-selectively onto a solid medium and subsequently checked for ICE presence by patching for tetracycline resistance. Tetracycline-resistant colonies were used as a template in a PCR reaction containing one arbitrary primer paired with an ICE-specific primer: oJ2260 (5’- GGCACGCGTC GACTAGTACN NNNNNNNNNT GATG) paired with either oJ2263 (5’- CTATTAGAAA GAAGATTAGC TGCAAACATC) for ICE*Bs1*/hybrids or oELC1009 (5’- GACATGCTAA TATAGCCATG ACG) for Tn*916*. Purified PCR products were amplified using oJ2262 (5’- GGCACGCGTC GACTAGTAC) and either oJ2265 (5’- ATCAGAACGA CCAAACAATG GT) for ICE*Bs1*/hybrids or oELC1011 (5’- GAACTATTAC GCACATGCAA C) for Tn*916*. Products were then sequenced with oJ2265 or oELC1011 and mapped to the appropriate sequenced genome. Genbank accession numbers for these species are: *B. subtilis* (CP007800) (Smith et al., 2014), *E. faecalis*, (ASDA00000000.1), *E. caccae* (JXKJ00000000) (Zhong et al., 2017), and *E. durans* (ASWM01000000).

## Acknowledgements

We thank Jennifer Brophy for providing pJAB403 that was used to generate D-alanine auxotrophs, Laurel Wright for generating the Δ*orf21* allele described here, and Monika Avello for designing primers used for excision assays.

Research reported here is based upon work supported, in part, by the National Institute of General Medical Sciences of the National Institutes of Health under award number R35 GM122538 to ADG. Any opinions, findings, and conclusions or recommendations expressed in this report are those of the authors and do not necessarily reflect the views of the National Institutes of Health.

---

**Table S1.**
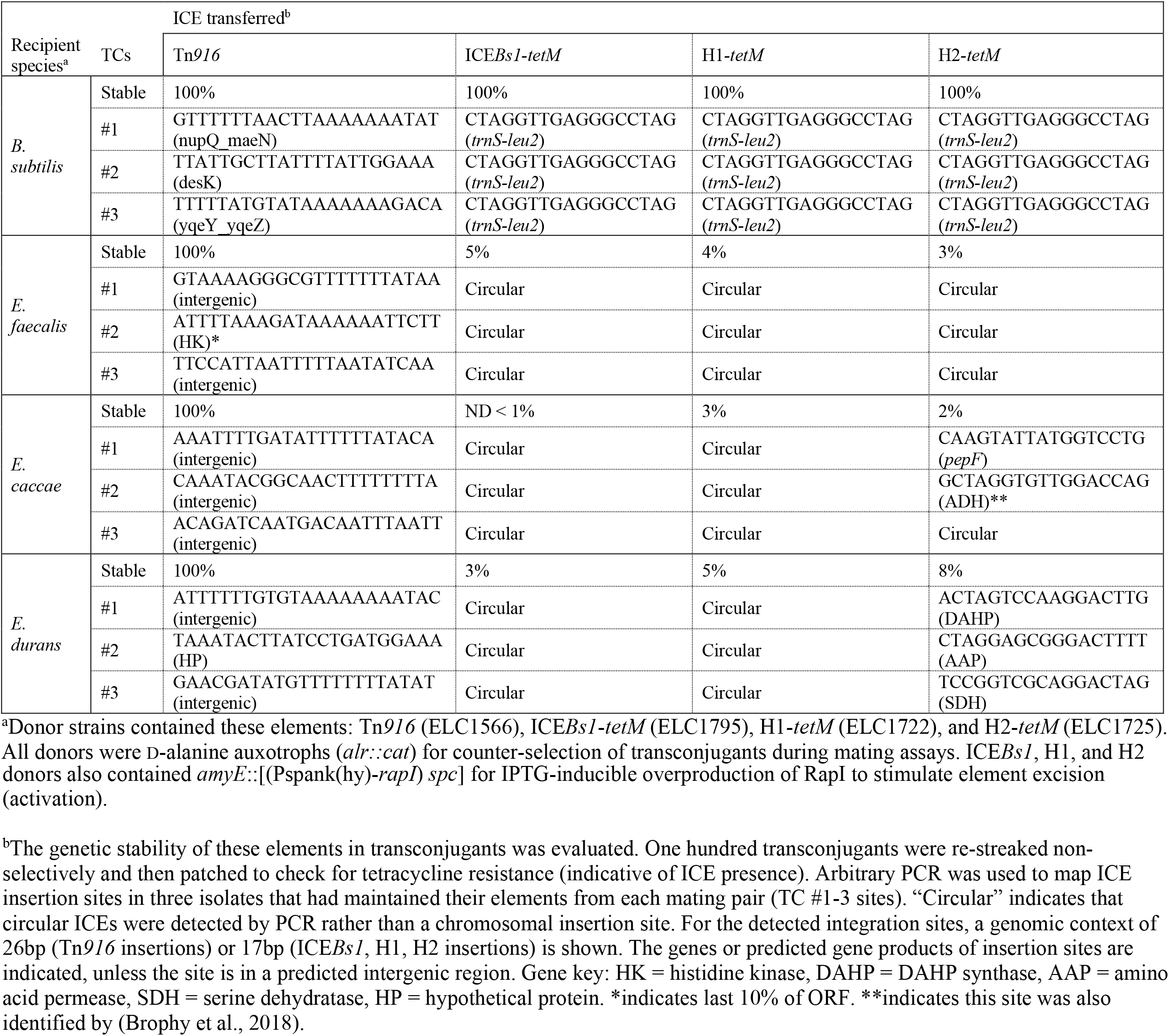
Mapping ICE integration sites in transconjugants.

